# Further studies of ion channels in the electroreceptor of the skate through deep sequencing, cloning and cross species comparisons

**DOI:** 10.1101/274449

**Authors:** William T. Clusin, Ting-Hsuan Wu, Ling-Fang Shi, Peter N. Kao

**Affiliations:** Department of Medicine Stanford University School of Medicine 300 Pasteur Drive Stanford Ca, 94305

## Abstract

Our comparative studies seek to understand the structure and function of ion channels in cartilaginous fish that can detect very low voltage gradients in seawater. The principal channels of the electroreceptor include a calcium activated K channel, whose α subunit is *Kcnma1,* a voltage-dependent calcium channel, *Cacna1d,* and a relatively uncharacterized K channel which interacts with the calcium channel to produce fast (20 Hz) oscillations. Large conductance calcium-activated K channels (BK) are comprised of four α subunits, encoded by *Kcnma1* and modulatory β subunits of the *Kcnmb* class. We recently cloned and published the skate *Kcnma1* gene and most of *Kcnmb4* derived from using purified mRNA of homogenized isolated electroreceptors. Bellono et al. have recently performed RNA sequencing (RNA-seq) on purified mRNA from skate electroreceptors and found several ion channels including *Kcnma1*. We searched the the Bellono et al RNA-seq repository for additional channels and subunits. Our most significant findings are the presence of two Shaker type voltage dependent potassium channel sequences which are grouped together as isoforms in the data repository. The larger of these is a skate ortholog of the voltage dependent fast potassium channel Kv1.1, which is expressed at appreciable levels and seems likely to explain the 20 Hz oscillations believed to occur in vivo. The second was more similar to Kv1.5 than to Kv1.1 but was somewhat atypical. We also found a beta subunit sequence (*Kcnab2*) which appears not to cause fast inactivation due to specific structural features. The new channels and subunits were verified by RT-PCR and the Kv1.1 sequence was confirmed by cloning. We also searched the RNA-seq repository for accessory subunits of the calcium activated potassium channel, *Kcnma1,* and found a computer generated assembly that contained a complete sequence of its beta subunit, *Kcnmb2.* Skate *Kcnmb2* has a total of 279 amino acids, with 51 novel amino acids at the N-terminus which may play a specific physiological role. This sequence was confirmed by PCR and cloning. However, skate *Kcnmb2* is expressed at low levels in the electroreceptor compared to *Kcnma1* and skate *Kcnmb1* (beta1) is absent. The evolutionary origin of the newly described channels and subunits was studied by aligning skate sequences with human sequences and those found in related fish: the whale shark (*R. typus*) an elasmobranch, and ghost shark (*C.milii*). There is also homology with the lamprey, which has electroreceptors. An evolutionary tree is presented. Further research should include focusing on the subcellular locations of these channels in the receptor cells, their gating behavior, and the effects of accessory subunits on gating.

## Introduction

Activation of potassium selective repolarizing ion channels by intracellular calcium is a fundamental principle of biology that was first appreciated in the mid 1970’s. In 1972, Meech [1] showed that injection of calcium ions into nerve cells with a microelectrode produced hyperpolarization that was due to an increase in potassium permeability. In 1974 he proposed several ways in which calcium ions might produce a generalized change in potassium permeability, which included the possibility of a “chemical reaction” with a site on the inner surface of the membrane [2]. But he eventually settled on the idea that excitable membranes contain a separate class of potassium selective channels that can be opened by intracellular calcium [3,4]. At about the same time, existence of distinct calcium activated potassium channels was proposed to explain the repolarization of excitable receptor cells in the electroreceptor of the skate (ampulla of Lorenzini) [5-10]. The action potential in these receptor cells was found to be mediated by an inward calcium current, and several lines of evidence showed that repolarization of this action potential could only occur when there was net calcium influx [6-8].

Verification that calcium activated potassium channels exist as distinct molecular entities came with the development of the patch clamp technique [11-12]. Subsequent work showed that calcium activated potassium channels are widespread, and that there are two principal types of calcium-activated potassium channels, large conductance (BK) and small conductance (SK) [13-14]. There is one type of BK channel, which is called *Kcnma1*, while there are four types of SK channels in vertebrates called SK1-4 (or *Kcnn1-4*). BK channels have a large conductance of 250-300 PS, while SK channels have a small conductance of 9-14 pS for SK1-3 and 10-42 pS for SK4. Large conductance calcium activated potassium channels are formed by a tetramer of alpha subunits, each of which may have accessory beta subunits [15-17]. Regulatory gamma subunits may be present as well. The alpha subunit of the BK channel has seven transmembrane regions (S0 to S6) and a large intracellular region which contains the molecular apparatus for sensing calcium and causing the conformational change which opens the channel. A conspicuous calcium binding site, called the calcium bowl, has four aspartate residues and is near the 3’ end of the alpha subunit. The calcium bowl is preceded by two regulatory complexes (RCK1 and RCK2) which exert mechanical force to open the channel by means of a linker that connects S6 to RCK1. A second calcium binding site (a single aspartate residue) is near the beginning of RCK1, 24 residues beyond the end of the S6-RCK1 linker. Both calcium binding sites participate in channel opening.

Most BK channels are sensitive to both intracellular calcium and voltage, and can be opened either by membrane depolarization or by a rise in intracellular calcium [18]. However the degree of sensitivity to voltage can vary, and in the skate electroreceptor, the calcium activated potassium current appears to be exclusively sensitive to calcium and insensitive to voltage [5-8]. The gating properties of the BK channels can be influenced by the type of beta subunits that are present [16, 18-21]. The beta subunits consist of two membrane spanning regions and contain fewer amino acids (about 200 as opposed to roughly 1100 for the alpha subunit). In mammals, there are four types of beta subunits called β1-β4.

The SK channel protein has six transmembrane spanning regions and has an intracellular region which is downstream from the membrane spanning regions [22-24]. Each channel has a tetrameric structure. SK channels are insensitive to voltage and do not have a voltage gate.

### Discovery of BK in the ampulla of Lorenzini through molecular cloning experiments

Rohman et al [25] initiated studies of the slo-1 gene that encodes the BK channel in brain tissue from teleost fish and in the skate, *L. erinacea.* They used degenerate primers that began with the highly conserved pore region, and were able to follow the sequence about 200 amino acids downstream. They showed that this region of the BK channel was highly conserved in many fish species. The proposal to obtain a complete BK sequence in the electroreceptor of *L. erinacea* was published as an abstract in 2010 [26]. Within a few months, the feasibility of doing this was greatly enhanced when a draft of the skate genome was obtained based on analysis of short reads of 50 to 100 nucleotides from a single late stage embryo [27]. As a first step, the skate version of *Kcnma1* was assembled from the library of skate genomic and transcriptome contigs by alignment with human *Kcnma1* [21]. This resulted in a full length nucleotide and amino acid sequence consisting of 1119 amino acids. Overall identity between the assembled skate and the human sequence was 96%, except that in the skate, a 59 amino acid region from the intracellular portion of the molecule was missing. The region that was missing is called the “strex” insert and its presence or absence is one of the most conspicuous features of alternative splicing of *Kcnma1*. Alternative splicing occurs at a total of seven specific sites in *Kcnma1* of vertebrate species [28]. After skate *Kcnma1* had been assembled, the resulting sequence was used to design PCR primers. These primers were used to amplify the full length sequence from purified mRNA that was obtained from 100 homogenized ampullae. The amplification product was ligated into a vector and further expanded in a bacterium. Two identical and complete sequences for skate *Kcnma1* were obtained based on the fact that there were two successful clones (21). The cDNA sequence for the alpha subunit was submitted to GenBank (accession number KJ756351.1). The corresponding amino acid sequence is AJP74816.1. A virtually identical amino acid sequence (97%) was very recently obtained and contributed to GenBank by Bellono, et al [29] (accession number AQV04218.1). Salient features of their sequence which agree with King et al. include the absence of the strex exon and the presence of exon 29 immediately before the calcium bowl. Bellono et al. showed that the alpha subunit of BK is the most abundant K channel in the ampulla, and is expressed at substantially higher levels (more than 35 fold) than the SK channels, *Kcnn2* and *Kcnn3*.

The physiological studies of the skate BK channel by Bellono et al reveal that the skate BK channel has a lower single channel conductance than mammalian or zebrafish channels due to specific amino acids in the S6 region (also called alternative splice site 2) which reduce the electrostatic interactions between potassium ions and the intracellular aspect (vestibule) of the channel pore. They show that the naturally occurring BK channel always has alternative exon 11 at splice site 2, whereas the other species noted above have exon 10. (Note that the “late stage embryo” form of BK alpha, described by King et al [21] also had exon 10 at splice site 2). To confirm their finding, Bellono et al made an artificial skate BK channel in which exon 11 was converted to exon 10 through point mutations that changed three charged amino acids. They found that the conductance of the artificial channel was similar to that of mouse wild type BK alpha. However, it is unclear if the data on the unmodified animal (n=7) is from more than one individual, and they did not report on juvenile (i.e. recently hatched) skates.

The Bellono, et al paper [29] also describes transcriptional profiling of the voltage dependent calcium channels (Cav) found in the ampulla of Lorenzini, along with physiological studies including heterologous expression. The pore forming subunit in the ampulla is Cav1.3, which is characteristic of hair cells in general. Skate Cav1.3 is 78% homologous to rat Cav1.3, but it is activated at more negative potentials, and shows reduced inactivation and a larger window current. This is consistent with voltage-clamp studies involving the dissected ampulla in which the early inward current, which we now know is carried by Cav1.3, does not inactivate during maintained depolarizations [6-8].

### Characterization of *Kcnmb4* in the ampulla of Lorenzini

While the paper of Bellono et al does not describe the beta subunits of *Kcnma1*, King et al were able to assemble a full length sequence for the β4 subunit (*Kcnmb4*) using the library of skate genomic and transcriptome contigs [21]. Appropriate primers for *Kcnmb4* were also designed and it was possible to amplify and then successfully clone two of the three exons of skate *Kcnmb4* (exons 1 and 3) and to also obtain a partial sequence for exon 2. However based on the Bellono data exression of *Kcnmb4* in the skate electroreceptor is low and it was not possible to complete the amino acid sequence using the deep sequencing data. In higher vertebrates such as mammals four distinct beta subunits designated as *Kcnmb1*-*4* have been found. The evolutionary origin of beta subunits is thought to have involved gene duplication.

### Evidence for a voltage-dependent potassium channel in the receptor cells

Physiological experiments suggest that the basal membranes of the receptor cells can give rise to oscillatory responses that involve an interaction between voltage dependent calcium channels and a potassium channel whose activation and deactivation are faster than that of the BK channel current produced by the apical faces [8-10]. These oscillatory responses can be abolished by bath application of known K channel blockers (tetraethylammonium [TEA] or strontium), which selectively affects the basal membranes. 20 Hz oscillatory responses could be observed during weak excitatory command pulses in voltage clamped ampullae [8-10]. Alternatively spontaneous 20 Hz oscillations of epithelial current occurred when physiological conditions were simulated by placing the canal in an air gap and short circuiting the preparation using a salt bridge [9]. Under these conditions, the spontaneous oscillations were accentuated when an excitatory voltage step was applied, or seen as an off-response (anode break) at the end of an inhibitory stimulus. Transmembrane oscillations of the basal surface of the receptor cells could be confirmed by recording the post-synaptic potential (PSP) of the afferent nerve in the presence of tetrodotoxin [8,9]. The 20 Hz oscillations in the PSP were exactly correlated with oscillations in the epithelial current [8]. It was suggested that the basal membranes contain voltage dependent potassium channels that explain these results, but the gating mechanism was not established with certainty, nor was any specific type of K channel implicated. Similar phenomena have been described in vertebrate neurons and in photoreceptors [Reviewed in ref. 10].

Modrell et al [30] have recently studied ion channels in the ampullary electroreceptors of the paddlefish which has strong similarities to skate electroreceptors. They found that the pore forming unit is Kv1.5 (*Kcna5*), which is a fast neuronal potassium channel of the type that would explain the results in skate electroreceptors. The paddlefish Kv1.5 channels also contain the beta subunit, *Kcnab3,* which is thought to cause fast inactivation of the channel during co-expression. Modrell et al suggest that non-teleost electroreceptor physiology may be may be conserved across jawed vertebrates, which would imply that the fast potassium channel in the basal membranes of skate electroreceptor cells is Kv1.5. Based on this work, we searched the deep sequencing data of Bellono et al to determine whether Kv 1.5 or another related channel in the “A current” (Shaker) family is significantly expressed in skate electroreceptors. In fact, we found that two Shaker type channels are expressed in the ampulla of Lorenzini, which could both be confirmed by PCR and cloning. One is a close homolog of Kv1.1 and the other has fewer amino acids and is similar to KV1.5. The data repository also contained a sequence for a beta2 subunit of Kcnma1, which was confirmed by PCR and cloning. A preliminary communication of some of these results has appeared [31].

## Materials and Methods

About 100 ampullae were dissected from each of two pairs of adult skates as described previously [21]. The skate ortholog of human *Kcnmb2* was identified by analysis of deep sequencing data recently obtained by Bellono et al, as described below. The deep sequencing data were derived from a single adult skate. The data of Bellono et al (NCBI GSE93582) contains computer generated nucleotide assemblies, and one particular assembly with unique similarity to human *Kcnmb2* was found by alignment. The nucleotide sequence number is c77966_g1. The nucleotide sequence predicted an open reading frame that was compared to *Kcmmb2* in other species.

Similar search techniques were used to find assemblies that encoded skate orthologs of Shaker type potassium channels and to find the open reading frame within those assemblies. This approach identified sequence c89985_g2 as having isoforms with strong homology to Kv1.1 and Kv1.5 as described below. A beta subunit, *Kcnab2*, was found using a similar strategy.

PCR was used to independently confirm the *Kcna1, Kcnab2* and *Kcnmb2* orthologs found in the deep sequencing data. Total RNA was isolated from the homogenized tissues using Trizol regent (Invitrogen, CA USA) following manufacturers protocol. cDNA was synthesized using SuperScript II Kit.™ First – Strand Synthesis System for RT-PCR (Invitrogen) using oligo (dT)_30_ primer following the manufacturers protocol. PCR primers were as follows. The primers for Kv1.1 (*Kcna1*) were Start/Plus: 5’ cggcgcgcctcactgaaggaagc 3’; End/Minus: 5’ gctctggccacggagcccgcag 3’. The internal primers used for cloning of Kv1.1 were Prox Third Plus 5’cgcgaggacgaaggcttcatc3’; Prox Third Minus 5’cacgcttcctgtcgtccttcagc3’; Distal Third Plus: 5’ccagcatgagggagctgggtcttc3’;Distal Third minus: 5’cgatcttacctccgatggtgaccgg3’ The primers for Kv1.5 (Kcna5) were Start/Plus: 5’gtccgcagacccgtcaacgtctccatcg3’; End/Minus: 5’ccttctatggataaaactaatctgtaatgttcaaacaatttgc. The primers for the beta subunit (*Kcnab2*) were Start/Plus: 5’ ccattcattcttgtggctgaagcagctgacctg 3’; End/Minus: 5’ gcttgagcaggcaaagaagtttgcttttgaaatg 3’. The four primers for *Kcnmb2* are given below.

For cloning experiments, the amplification product was ligated into a PCR2.1 vector (Invitrogen) and sequenced using an Applied Biosystems 3100 DNA sequencer (Applied Biosystems, Foster City, CA, USA). By finding the open reading frames, it was possible to identify the amino acids in both the deep sequencing assembly and the PCR amplification product obtained from the purified mRNA. The computer program SkateBase (www.skatebase.org) was used to confirm the presence of the *Kcnmb2* sequence in the skate genome and to identify the specific genomic contigs. In general, the discrepancies between the deep sequencing assemblies and the cloning results are minor, reflecting rare amino acid variations between individual skates.

## Results

### Identification of skate Kv1.1 in the deep sequencing data

As noted in the introduction, it is likely that the skate electroreceptor has a voltage dependent potassium channel located in the basal membranes of the receptor cells. Because of the importance of this channel it seemed likely that a computer generated assembly that contained this potassium channel would be found in the data deposited by Bellono et al in NCBI GEO. Based on belief that this might be Kv1.5, a search for alignments against a human Kv1.5 sequence with the Bellono deep sequencing data was undertaken. This search pointed to assembly c89985_g2, which is represented by eight assemblies (isoforms) designated as i1 – i8. Isoform 1 was analyzed with a DNA to protein converter tool, which revealed a large open reading frame. The protein translation of that isoform is shown in Fig. 1. Two of the other isoforms, i2 and i5 give exactly the same protein which contains 490 amino acids and ends with a stop codon (*). When a cross species search for this protein was performed using Blastp, it was found to have much stronger similarity to human Kv1.1 (NP_000208.2) and other Kv1.1 proteins than to Kv1.5. (Kv1.1 is a Shaker-related potassium channel that in humans is encoded by the *Kcna1* gene.) Alignment of skate Kv 1.1 with human Kv1.1 showed 86% identities (Fig. 2), while alignment of the skate sequence with human Kv1.5 (AAH99666) gave 65% identities The strongest alignment for skate Kv1.1 is found in the whale shark, *R.typus*, (XP_020390878.1) as illustrated in Fig.3. The alignment shows 96% identities and both proteins have 490 amino acids. As stated above, the PCR product was ligated into a cloning vector and the cloned product was sequenced to confirm its identity with the deep sequencing assembly. 484 of the 490 amino acids were identical in the cloned PCR product and the Bellono assembly. There were six sporadic single amino acid changes shown as red letters in Fig. 1. Expression of mRNA for skate Kv1.1 was further demonstrated by PCR which produced the expected PCR product containing 1470 bases. The expression level estimated from the transcriptome data of Bellono et al. for skate Shaker channels (Kv1.1 plus Kv1.5) and was substantially higher in the ampulla than the canals, skin or liver, and was 8.3% of the value obtained for *Kcnma1* (Table 1).

**Table 1.** Expression levels of Kcnma1, Kcnmb2, Kcnmb4, Kv1.1, and Kvβ

|  | Ampullae | Ampullary canals | Skin with no canals | Liver |
| --- | --- | --- | --- | --- |
| Kcnma1 | 694.29 | 5.16 | 50.25 | 2.06 |
| Beta2 | 0.62 | 3.16 | 0.0 | 0.14 |
| Beta4 | 0.56 | 0.0 | 0.0 | 0.0 |
| Kv1.1+ 1.5 | 57.49 | 13.97 | 2.79 | 34.32 |
| Kvβ2 | 380.00 | 148.00 | 196.00 | 111.00 |

Levels calculated from GSE2455279 by RSEM (RNA-Seq by Expectation Maximization) Four samples.

**Fig 1.**
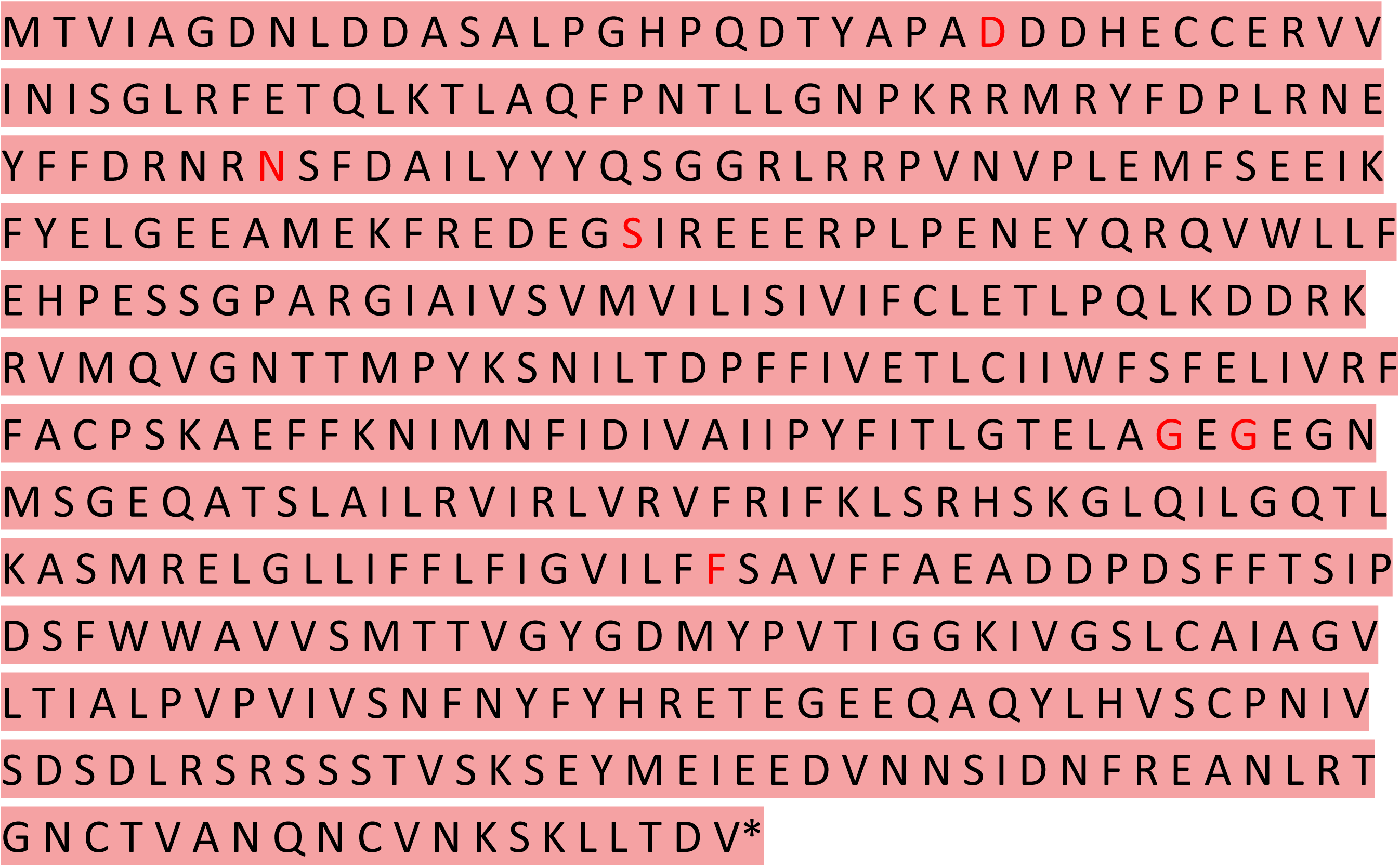
Sequence of skate Kv1.1 found as a deep sequencing assembly contributed to a repository by Bellono et al. [29.] The sequence contains 490 amino acids and is well aligned with the human and whale shark orthologs. * marks the stop codon. This sequence is specified by assemblies c89985_ g2, i1, i2 and i5. Besides the presence of a start methionine and a stop codon, the sequence is identifiable as an ion channel by the presence of a “pore region” which contains four closely spaced arginine residues. These are in the 8^th^ line of the figure and contain the letters “SLAILRVIRLVRVFR.” Red letters indicate six amino acids that were different in the cloned sequence than the assembled sequence. The six amino acid changes were N to D, R to N, F to S, E to G, D to G and S to F, where the red letter shows the amino acid obtained by cloning for each pair.

**Fig 2.**
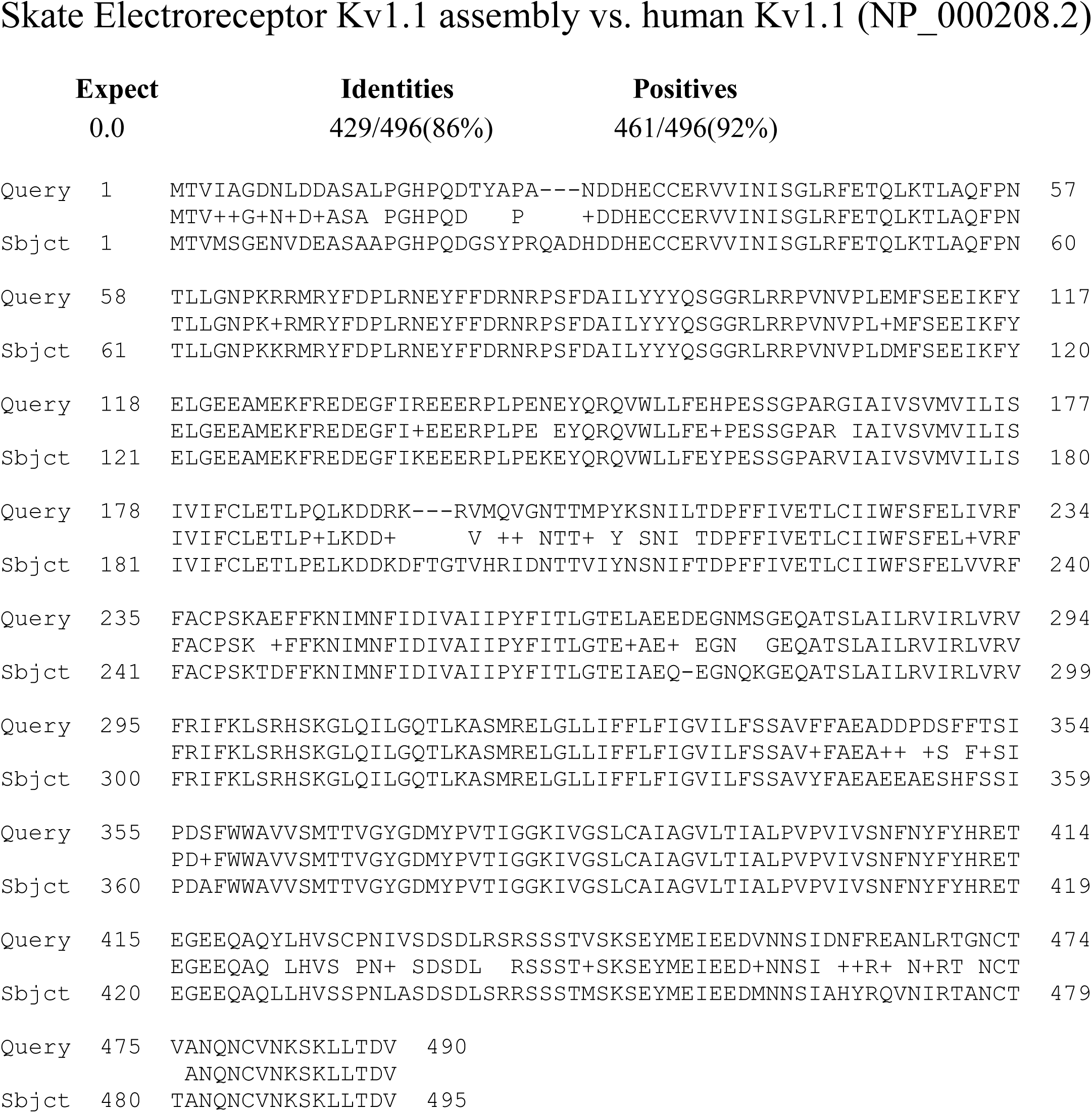
Alignment of the protein sequence of skate Kv1.1 (Query) with the human isoform (Subject). 86% of the amino acids are identities. This alignment is superior to the alignment with any of the other seven human Kv1 sequences.

**Fig 3.**
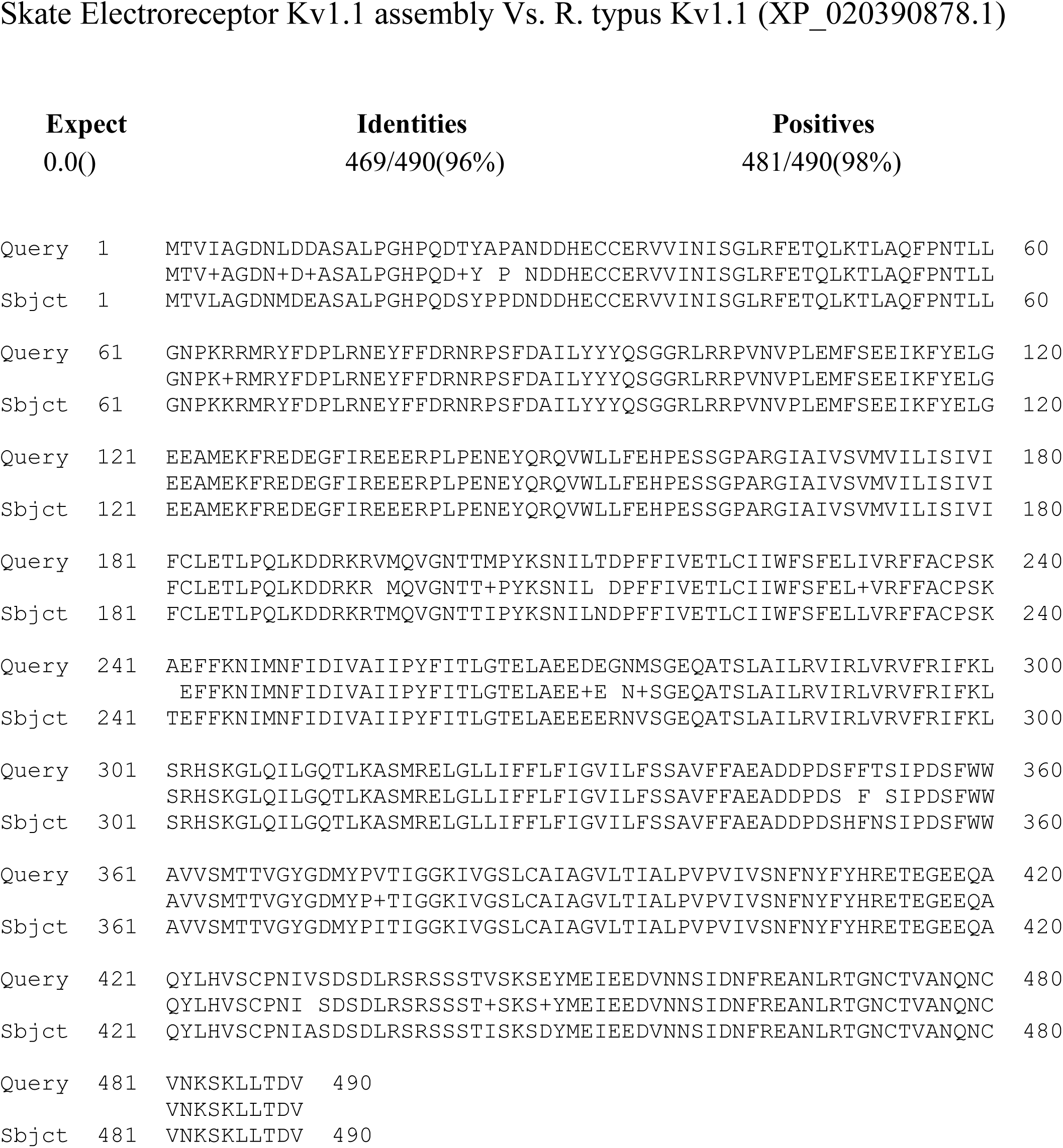
Alignment of the protein sequence of skate Kv1.1 to the ortholog found in *R. typus*, the whale shark, which is an elasmobranch. Both sequences have 490 amino acids and there are 96% identities.

The Kv1 family of potassium channels consists of eight members called Kv1.1-Kv1.8, and is also called the shaker potassium family or family A [35]. It is not clear if eight separate Kv1 genes are present in the skate, and the present paper considers only the electroreceptor. However there are complete sequences available for the human isoforms. When skate Kv1.1 was compared against the other seven human members of the Kv1 family, the strongest alignment was against human Kv1.2 (NP_004965.1) where 81% identities were found (E=0) and the proteins were identical at the 5’ and 3’ ends. Human Kv 1.2 contains 499 amino acids. Kv1.1 channels are the most prevalent potassium channels in the nervous system of animals and are highly expressed in both axons and cell bodies [36].

Kv channels are ubiquitous among animals and are significantly conserved in non-vertebrate species, indicating that they were present before vertebrates evolved. To verify this, the sequence of Kv1.1 of the whale shark, *R. typus* was aligned against the Kv1 protein of the lamprey as shown in Fig. 4 (lamprey sequences in dark red). The lamprey Kv1 sequence was curated as KV1.7 (Uniprot S4RCX3) although only two lamprey Kv channels have been curated. The lamprey Kv protein has 331 amino acids, and the skate form has 99 unaligned AAs at the 5’ end and 79 at the 3’ end. Overall alignment is highly significant (6E-141) and is most conspicuous near the lamprey 3’ end.

**Fig 4.**
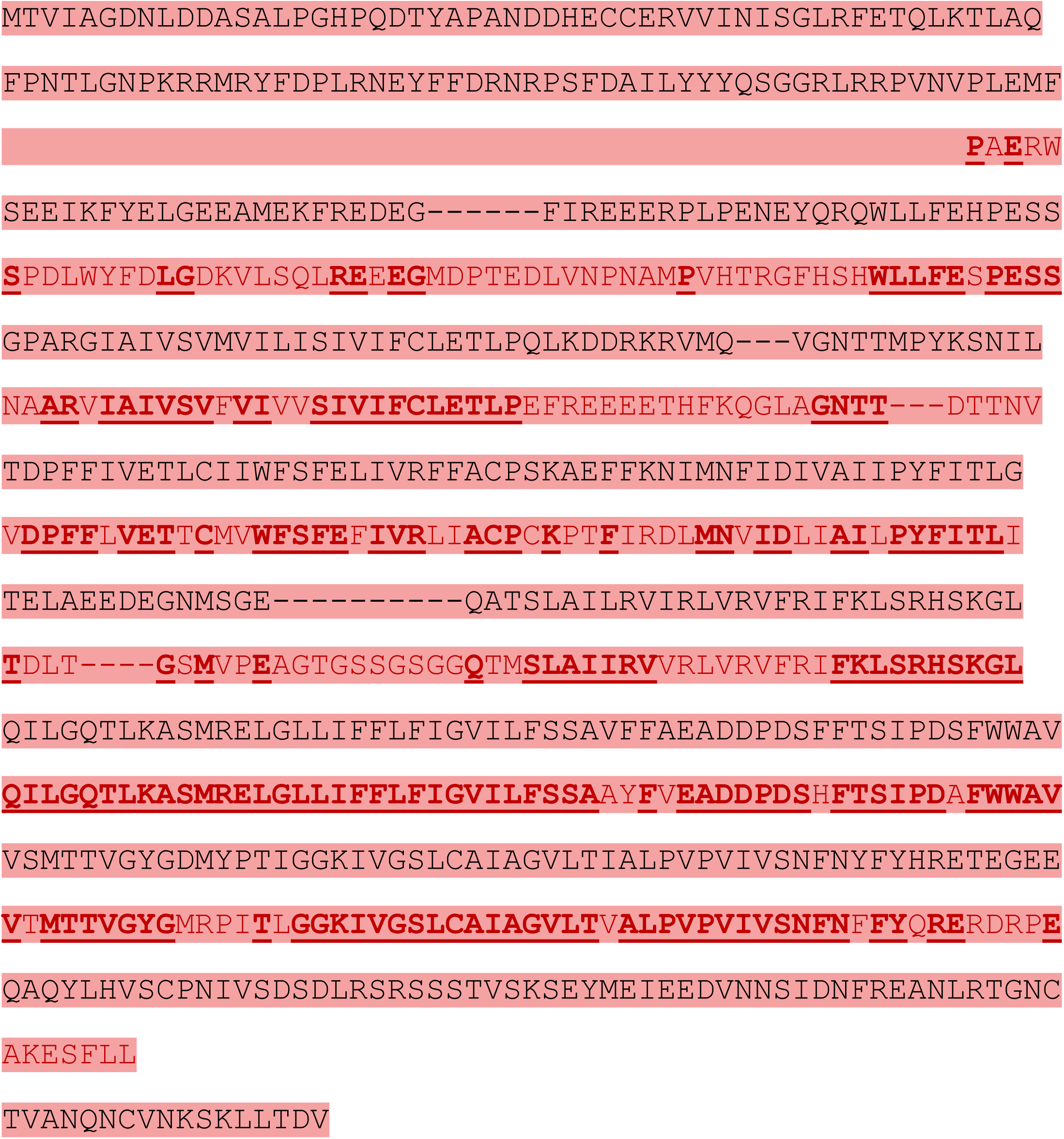
Alignment of the protein sequence of Kv1.1 in the skate electroreceptor (black letters) with the ortholog of Kv1.1 found in the lamprey genome. (red letters). Lampreys have electroreceptors called end buds. Although the Kv1.1 gene expressed in the lamprey has fewer amino acids than the skate, the alignment is highly significant.

### Identification of skate Kv1.5 (short Kv1.5) in the deep sequencing data

Three other isoforms found in assembly c89985_g2, numbered i6, i7, and i8, have an open reading frame which codes for the alpha subunit of a second Shaker type voltage dependent potassium channel, whose amino acid sequence is shown in Fig. 5. The pore region described above, which has four charged arginines, is also present in Kv1.5. A comparison of the Kv1.1 and Kv1.5 sequences show that there are 66 downstream amino acids (starting with SLAILRV..) which are identical in the two sequences. The 377 amino acids in skate Kv1.5 (Kcna5) are coded for by 1131 nucleotides given in the deep sequencing assembly. Appropriate PCR start and end primers for Kv1.5 were designed, as stated in the methods. These primers gave a PCR product of appropriate size on polyacrylamide gel. This was repeated five times. The PCR product was sequenced and 374 of the 377 amino acids (99%) were confirmed. Those that were not confirmed are shown by red letters in Fig. 5.

**Fig 5.**
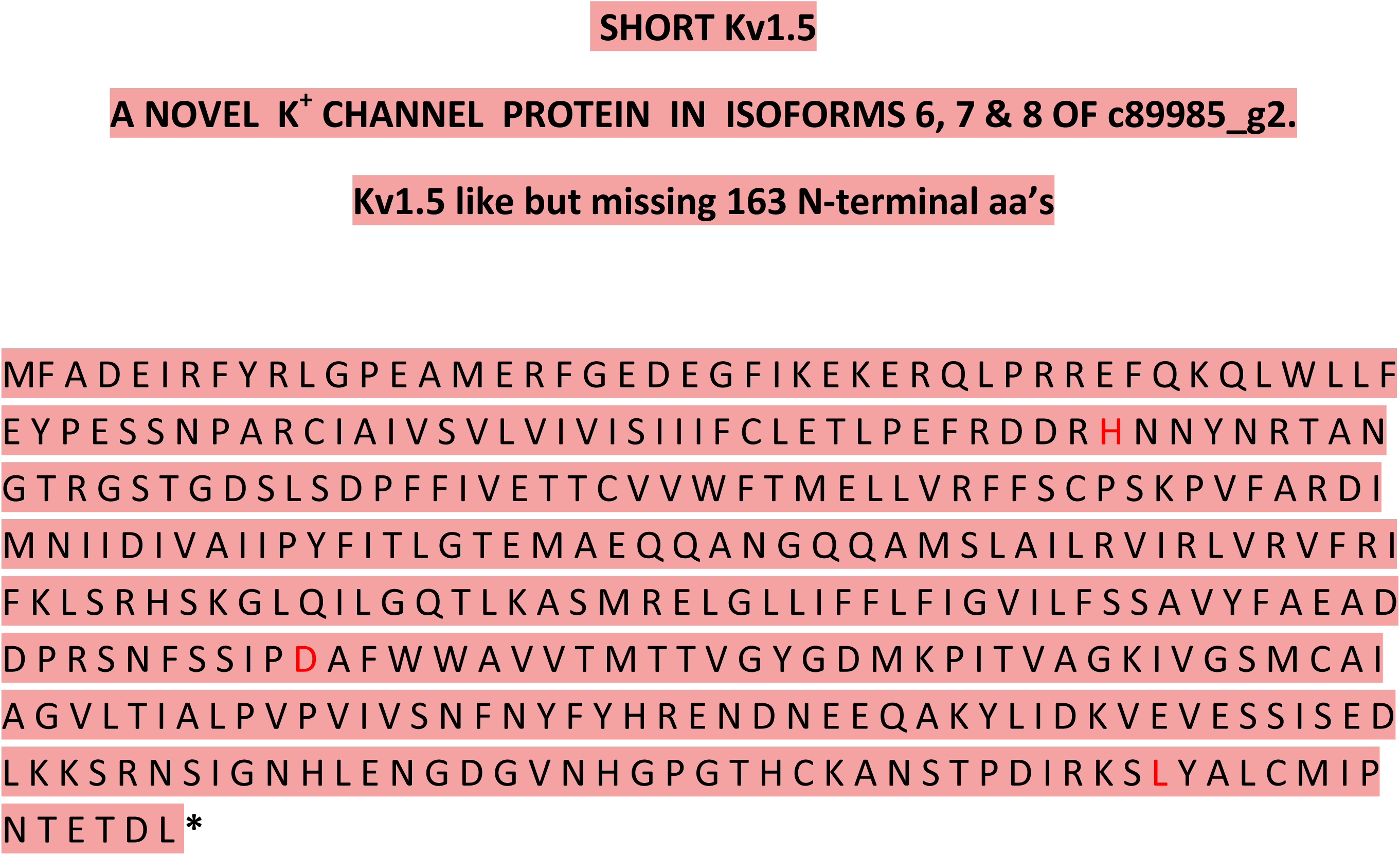
Amino acid sequence of Kv1.5 found in isoforms 6,7, and 8 of the deep sequencing data of Bellono et al (29). There are a total of 377 amino acids beginning with the start methionine. This sequence shows a high degree of alignment (E=0) with Kv1.5 of the whale shark, *R.* typus, (XP_020390867) except that the first 163 amino acids of the whale shark sequence are missing. There is complete alignment with skate Kv1.1 in the pore region and for 66 downstream amino acids. The presence of skate Kv1.5 in purified m-RNA from dissected ampullae was confirmed by PCR, and the PCR product was sequenced. Only three of the 377 amino acids could not be confirmed by sequencing the PCR product (red letters).

### Beta subunit of skate Kv1.1

Kvα subunits are integral membrane proteins that assemble as tetramers and can function as channels. Kvα can co-assemble with Kvβ subunits which are cytoplasmic proteins that associate with distinct cytoplasmic domains of Kvα. Modrell et al [30] reported that Kv1.5 is co-expressed with the beta subunit *Kcnab3* in the ampullary organs of paddlefish. Leicher et al [39] report that co-expression of human Kv1.5 with *Kcnab3* may produce channels that exhibit rapid inactivation, but this depends on the expression system that is used. Channels in the Kv1 family are known to associate with beta subunits designated *Kcnab1*. Based on this information, a search of the ampulla deep sequencing data was performed using blast P with human *Kcnab1* protein as the query. The search revealed four assemblies with a high degree of alignment which were c7914_g1 isoforms 1-4. Isoform 1 and 3 are identical and code for 353 AA’s while isoforms 2 and 4 are identical and code for 367 AA’s. When the sequences are aligned they are identical except that the shorter form has a characteristic gap of 14 AA’a compared to the long form.

When the above assemblies are searched for in the NCBI protein library using Blastp, they are consistently identified as coding for *Kcnab2* and not *Kcnab1*. This was true for all four isoforms of c79174_g1. The closest alignment was for *Kcnab2* in the ghost shark, *c. millii* (XP_007896081.1) and all listed alignments for other species were against *Kcmab2*. The protein sequence (Fig. 6) should therefore be designated as skate *Kcnab2* (also called Kvβ2). Expression of mRNA for skate *Kcnab2* was further confirmed by PCR, which produced the expected PCR product (1101 bases). The PCR for *Kcnab2* was repeated four times (primers given in Methods). The PCR product was sequenced, which showed that 94% of the amino acids were identical to those found in the deep sequencing isoforms.

**Fig 6.**
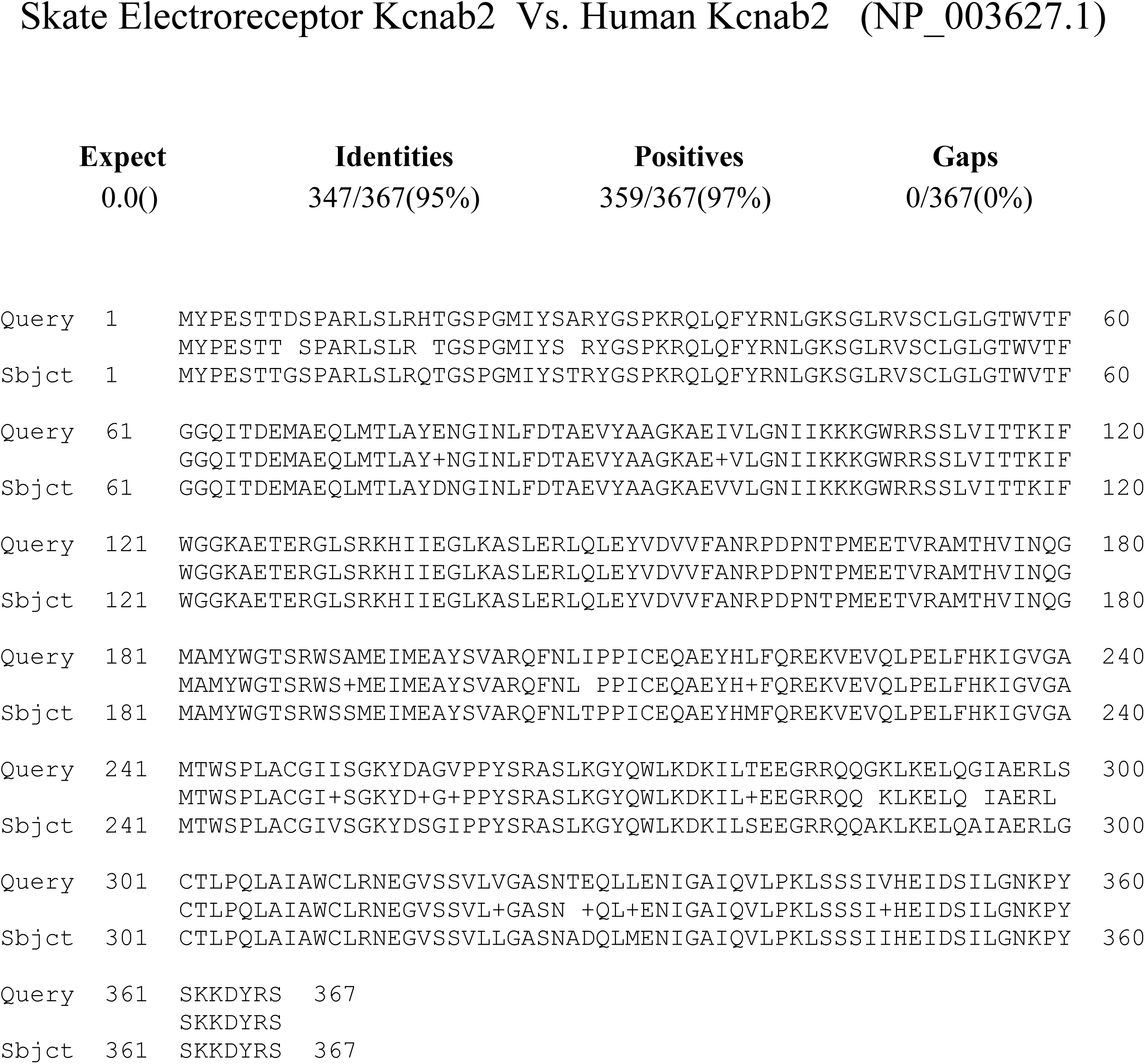
Assembly the skate *Kcnab2* obtained from c79174_g2 (isoform 1) of the deep sequencing file (Query) aligned against human Kcnab2 (NP_003627.1) (Subject). The sequences are highly conserved showing 95% identities. Two complete copies of this full length alignment were found in the deep sequencing file. Skate Kcnmb2 also aligns against human Kcnmb1, although the first 76 amino acids of the human form are missing and have no skate counterpart.

A detailed crystal structure has been published for the beta subunit of Kv1.1 in the rat, and its point of contact with the intracellular portion of the first membrane spanning region of the alpha subunit (40). Fig. 1B of cthis paper shows 10 specific numbered amino acids for the alpha subunit and 8 for the beta subunit all of which exactly correspond to the sequences for the skate subunits shown in Fig. 1 and 6.

An important feature of skate *Kcnab2* is that when it is aligned with the sequence of human beta1 (Q14722.1) the first 76 amino acids of the human sequence do not appear in the alignment, which starts with amino acid 77. Since the missing amino acids mediate ball and chain inactivation, it can be concluded that skate *Kcnab2* may not have this property.

### Discovery of skate *Kcnmb2* in the ampulla of Lorenzini through deep sequencing

Because of the usual importance of beta subunits in modulating the gating behavior of *Kcnma1*, the deep sequencing data deposited by Bellono et al in NCBI GEO was searched for computer generated assemblies that might contain the sequence of a beta subunit. An initial attempt was made to identify a skate version of *Kcnmb1* by alignment with the human ortholog. This lead to the discovery of a 940 base assembly (Fig. 7) which mostly consists of a single open reading frame which codes for *Kcnmb2* amino acids. While this assembly was found because of its strong alignment with human *Kcnmb1* (E=5e-32), the assembly was also checked for alignment against human Kcnmb2, which was even better (E=2e-84). For this reason, the assembly described above has been designated as “skate *Kcnmb2*.”

**Fig 7.**
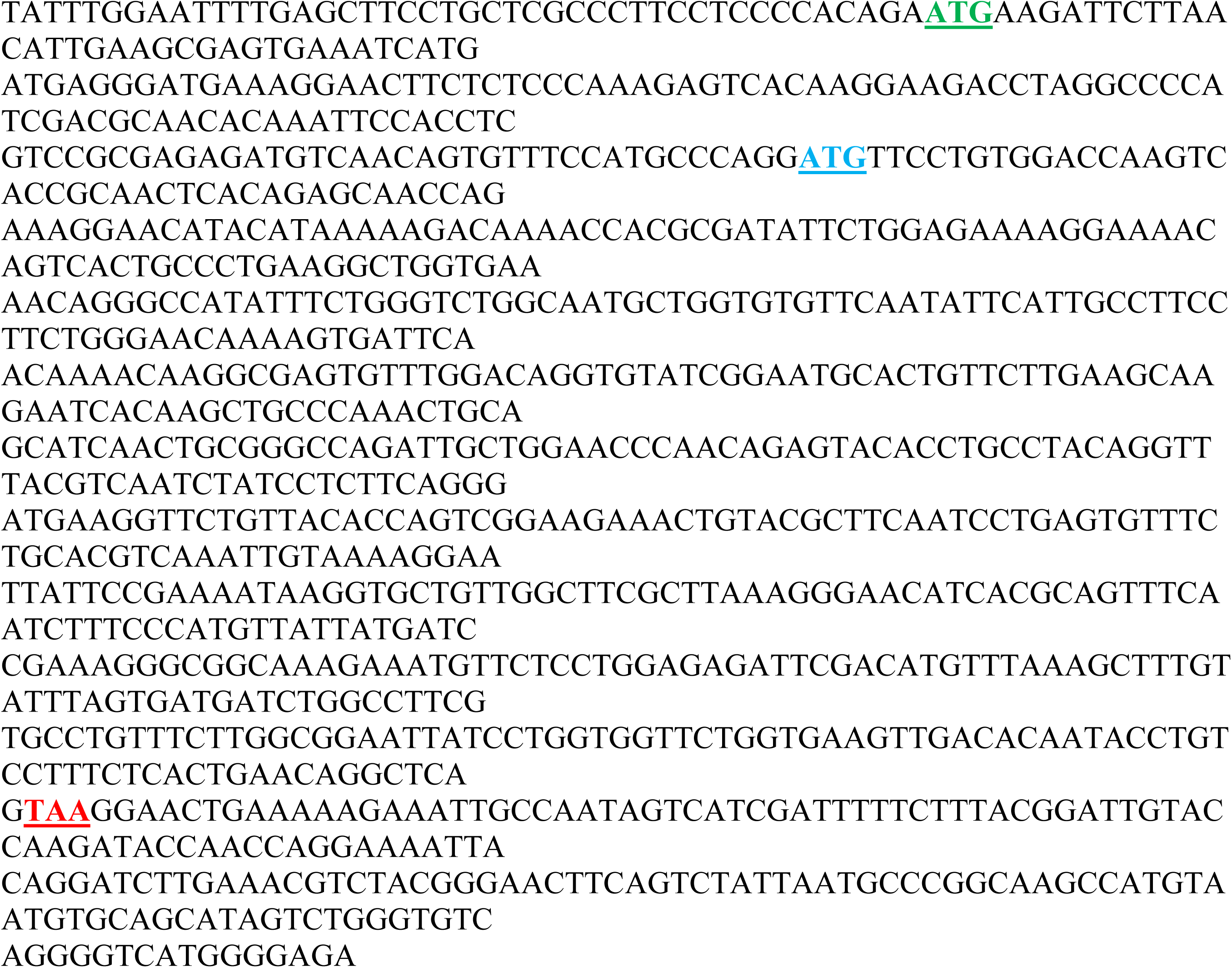
Nucleotide assembly which codes for the β2 subunit of the BK channel of the skate electroreceptor. The assembly was found by a search of deep sequencing data from purified mRNA from the skate electroreceptor. Data was obtained by Bellono et al [29] and deposited in NCBI GEO, accession number GSE93582; sample GSM2455279; assembly number c_7796_g1. The assembly includes of a 44 nucleotide 5’ untranslated region and a 170 nucleotide untranslated region after the stop codon (TAA, red). The first ATG (green) indicates the start methionine for skate β2, while the blue ATG indicates the consensus methionine for several species (see Fig. 10).

**Fig 8.**
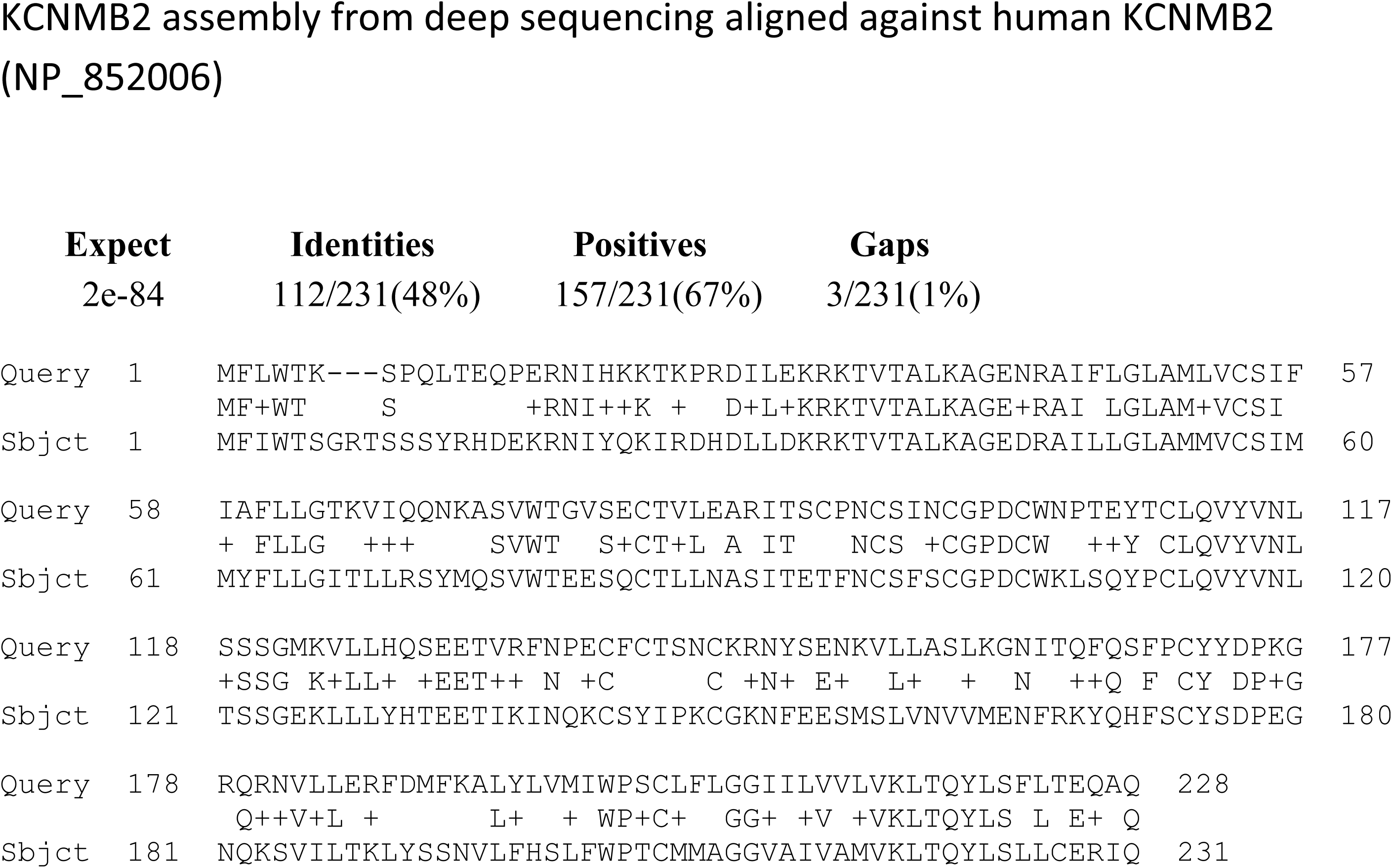
Skate *Kcnmb2* amino acid assembly obtained from deep sequencing of the electreceptor aligned against human *Kcnmb2*. The human sequence is the query. The subject amino acids in Fig. 8 are also shown in Fig. 9 beginning with aa 52.

A complete translation of the nucleic acid sequence in Fig. 7 is given in Fig. 9 and 10, which represents the full extent of skate *Kcnmb2*. Skate *Kcnmb2* contains 279 aa’s. The most noticeable distinction between skate and human *Kcnmb2* is that the skate ortholog has 51 aa’s at the beginning of the sequence (starting with MKILN.… and ending at MFLW…) which are not present in the human sequence.

**Fig 9.**
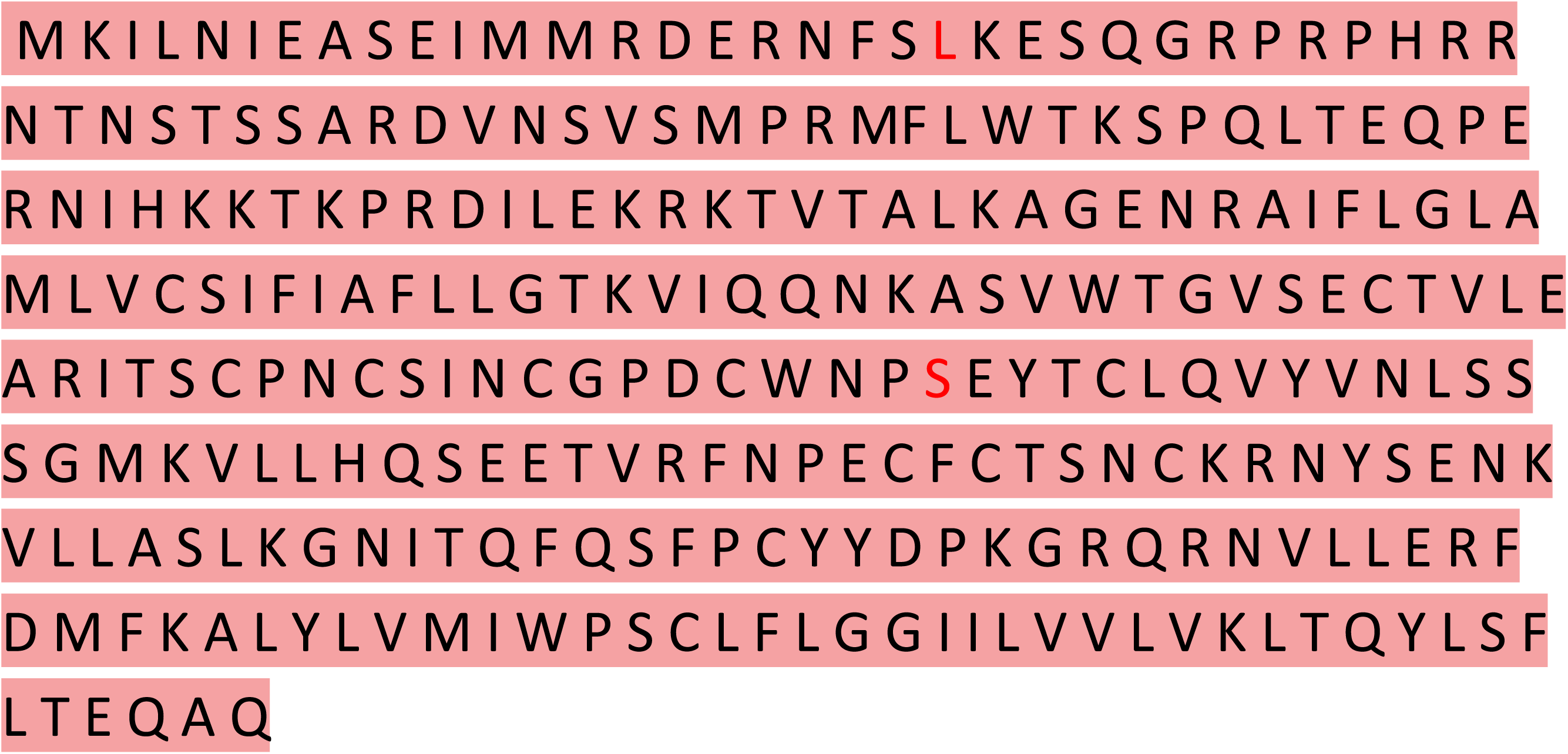
Amino acid sequence for skate *Kcnmb2* derived from the nucleotide assembly in Fig. 7. It contains 279 amino acids. Using the successful PCR primers, it was possible to confirm the amino acid sequence, to the extent that it was identical, and identify two amino acids (red letters) that were different in the cloned sequence. There were three single nucleotide polymorphisms in the cloned sequence as compared to Fig. 7. One of these (the middle of the three) was synonymous and two produced the amino acid changes shown.

**Fig 10.**
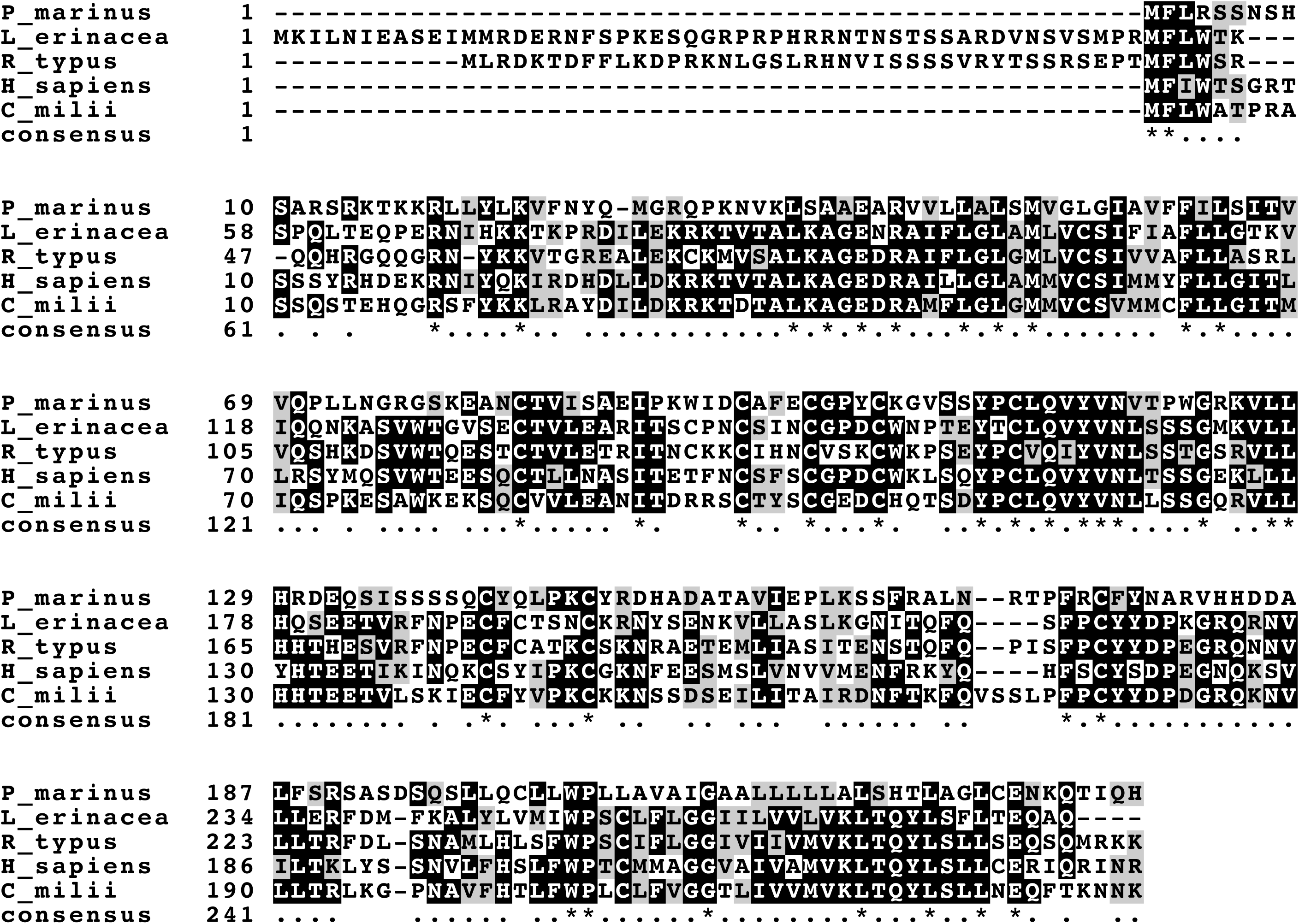
Alignment chart comparing *Kcnmb2* sequences in four species, the sea lamprey (*P. marinus)*, the skate (*L. erinacea*), the whale shark (*R. typus)*, the human, and the Australian ghost shark (*C. mili*). Both elasmobranchs, skate and *R. typus*, show a substantial number of amino acids prior to the consensus methionine (51 for *L. erinacea,* 40 for *R. typus*). Beginning with the consensus methionine, the first 45 amino acids of human and mouse β2 comprise the “ball and chain” motif, which is responsible for rapid inactivation. Rapid inactivation of BK channels has not been observed in the skate and might not occur, particularly if the initial 52 amino acids function as an “N-type inactivation-prevention (NIP).

### Comparison of Skate *KCNMB2* with other species

Fig. 10 illustrates alignment of *Kcnmb2* in four species; the lamprey, *P. marinus, L. erinacea, R. typus, Homo sapiens,* and *C. milii*. Black boxes show identities, grey boxes show similarities, and white boxes show amino acids unrelated to the others in a column. As noted above, three of the sequences begin with a “consensus” methionine and are approximately the same length, and the two elasmobranchs have amino acids prior to that, which begin with a methionine. It is possible that the additional amino acids have the function of disabling the “ball and chain inactivation mechanism (called N-type inactivation) that is usually present in BK channels that have a β2. Naturally occurring variants of the voltage dependent K_v_1.6 (which normally has N-type inactivation) have been described in which there are 42 additional N-terminal amino acids which serve as an “N-type inactivation prevention (NIP) domain” [32]. It is therefore possible that both *L. erinacea* and *R. typus* have naturally occurring β2 subunits that do not produce inactivation because inactivation is disabled by the NIP domain. This would be consistent with the electrophysiological data for *L. erinacea* [6-10].

In the case of K_v_1.6 a small negatively charged sequence (EFQEAE) appeared to be important for function [32]. Substitution of alanine, or lysine for the three glutamic acid residues (E27/30/32) conferred inactivation behavior. It is therefore possible that the two closely spaced glutamic acids in the N-terminal region of skate β2 (EASE in positions 7-10) play a similar role.

The hypothesis that the nucleotide assembly shown in Fig. 7 encodes “skate *Kcnmb2*” is even more strongly supported by using the program BLASTP to identify species where the sequence of *Kcnmb*2 is most similar to the skate. The strongest similarity to *L. erinacea Kcnmb2* is to the whale shark, *Rincodon typus,* where the extent of alignment in *Kcnmb2* is 268 amino acids, and the E value is 8e-99 (Fig 10 and 11). The whale shark *Kcnmb2* has 40 amino acids preceding MFLW and this region shows significant alignment with *L. erinacea* (13/40 identities). Second on the list of species where *Kcnmb2* is most closely aligned with *L. e*rinacea is the Australian ghost shark, *Chilorhinchus milii* (Uniprot V9KI 35, Fig. 10 and 12). The E value for this alignment is 1e-88 (125/232 identities). While *L. erinacea* and *R. typus* are both elasmobranchs, *C. milii* is a chimera which is a cartilaginous fish and has electroreceptors but belongs to the Holocephali subclass rather than elasmobranchii. The similarities in the sequence of *Kcnmb2* as quantified by the alignment data are therefore in line with the evolutionary relationship among the species studied.

**Fig 11.**
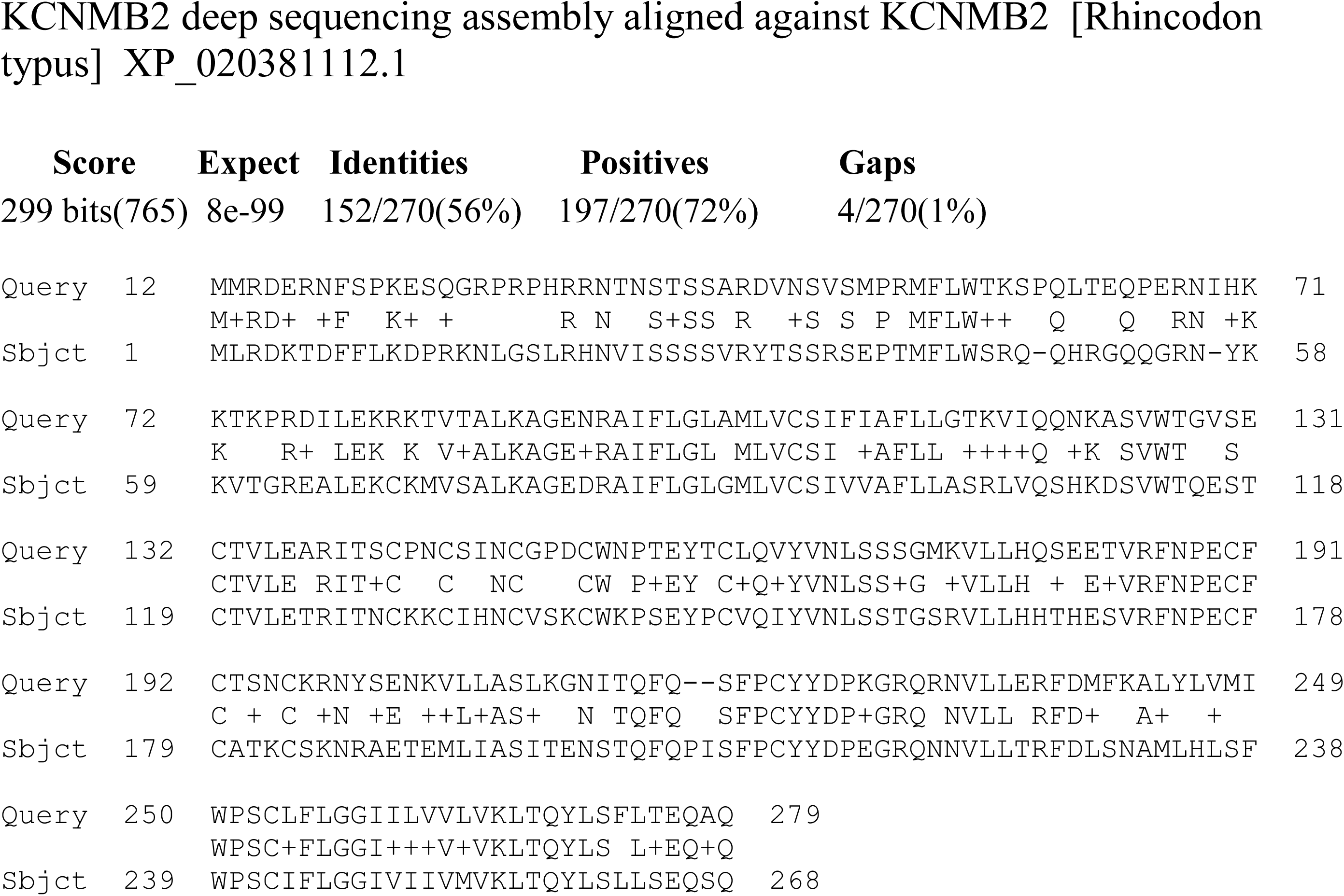
Alignment of amino acids between *Kcnmb2* (β2) in the skate electroreceptor (Query) and in the genomic sequences of *R. typus* (subject). Tissue source for *R. typus* was the spleen. Because the number of amino acids that precede the start methionine is less in *R.* typus, the alignment starts at the 12^th^ amino acid in *L. erinacea,* which is also a methionine. Alignment was performed with Blastp, and gave an e-value of 8e-99.

**Fig 12.**
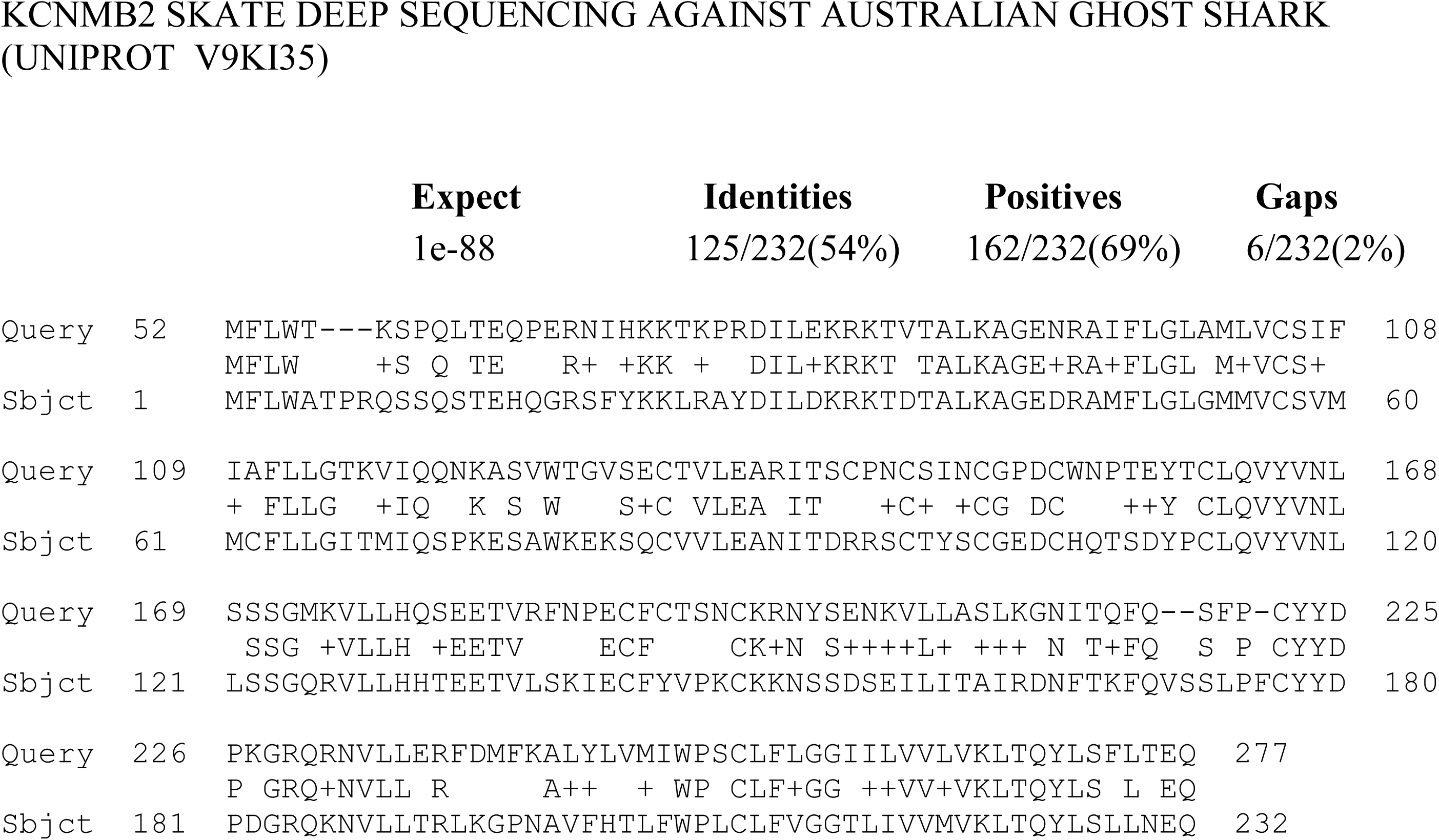
Alignment of amino acids between *Kcnmb2* of skate electroreceptors (Query) and in the Australian ghost shark, *C. milii,* as predicted from the genomic sequence. Neither sequence has amino acids prior to the consensus start methionine. Comparison of the sequences with BlastP gives a significance of 1e-88. *C. milii* is a chimera which has electroreceptors but is slightly less closely related than the whale shark, and is believed to have evolved earlier than elasmobranchs.

### Confirmation of the Skate *Kcnmb2* Sequence by PCR and Cloning

While the presence of a human β2 analog in the deep sequencing data from the ampulla is strong evidence that the gene for *Kcnmb2* is transcribed in the skate, this conclusion has been further confirmed by designing PCR primers from the nucleic acid sequence in Fig. 7, and then cloning the PCR product in E. Coli. The five PCR primers, three forward and two reverse, that were constructed for skate *Kcnmb2* are shown in Table 2. The start primer begins with the start methionine for skate *Kcnmb2*, which is indicated in green letters in Figure 7, while the end primer (2-) ends at the stop codon. Four specific pairs of PCR primers that successfully gave a PCR product of appropriate size. The primer pair consisting of 1+-2-gave a full length sequence of 848 amino acids. The other primer pairs tested were 1+1-(585 aa), 2+-1-(455 aa) and 3+2-(395 aa).

**Table 2.**
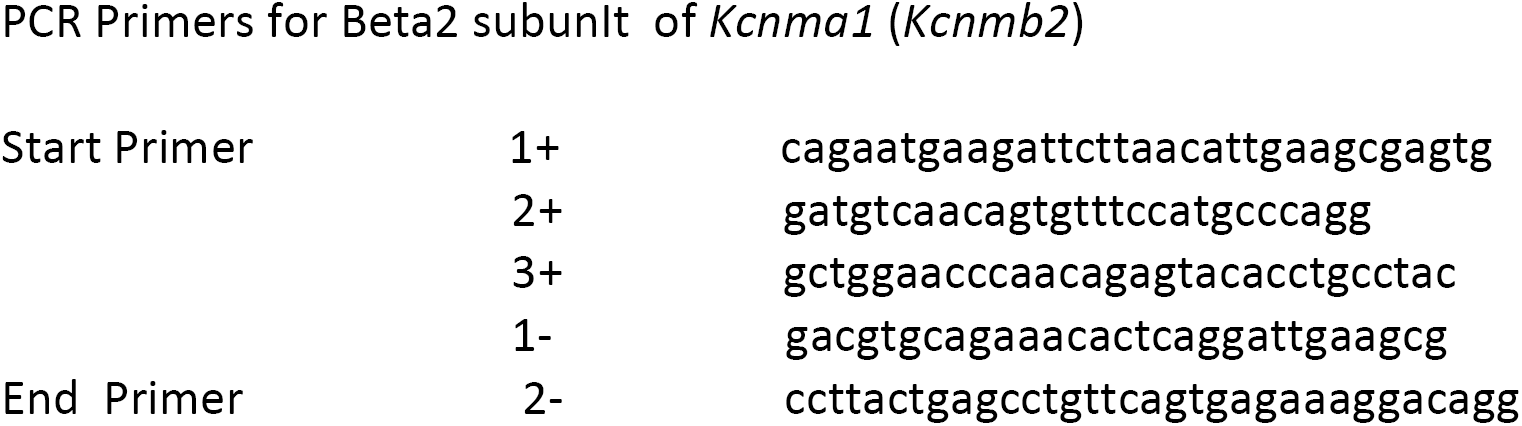
PCR Primers for Beta2 subunIt of *Kcnma1* (*Kcnmb2*)

The sequence of skate *Kcnmb2* in Fig. 7 was confirmed by cloning. The stop codon, TAA, is highlighted in red. The nucleotide sequence obtained from the successful clones (which was flanked by vector sequences) included a brief 5’ untranslated region and a longer 3’ untranslated region. The confirmed coding sequence began with the start methionine, ATG, which is shown in bold green in Fig. 7, and ended with the nucleotides specified by the 2-primer. The nucleotide sequences contained in the cloned PCR product and the deep sequencing assembly were identical except for three single nucleotide polymorphisms (SNP’s), one of which was a synonymous SNP and two of which changed a single amino acid. The two amino acid changes (shown in red in Fig. 9) are P to L at amino acid number 21 and T to S at amino acid 157. It is seen that sequence of *Kcnmb2* is highly similar among individual skates.

### Relative expression levels of Kcnma1, Kcnmb2 and Kcnmb4

An important caveat with respect to the interpretation of data on *Kcnmb2* and *Kcnmb4* is their low expression levels with respect to *Kcnma1* (see Table 1). For *Kcnmb2* the expression level is approximately 1000 times lower than *Kcnma1* and 10,000 times lower for *Kcnmb4*. While beta subunits of the BK channel are known to be present in other organs, such as the brain and smooth muscle, the data in Table 1 together with the absence of detectable β1 in the deep sequencing repository suggest that the BK channels in the electroreceptor cells have no beta subunits at all. Many expression studies including single channel recording have been obtained with *Kcnma1* channels that did not have beta subunits, and a mouse model exists in which *Kcnma2-/-* animals were created and survived [19]. Thus the results in Table 1 are not inconsistent with existing knowledge. A complicating factor is that a substantial proportion of *Kcnma1* molecules reside in intracellular compartments including the mitochon-dria [33], nuclear envelope [34] and sometimes the nucleoplasm. Further studies are needed to investigate the localization of *Kcnma1* transcripts in the ampulla of Lorenzini and the role of beta subunits.

## Discussion

The objective of this research has been to obtain additional information about ion channels that are expressed in the ampulla of Lorenzini of the skate using a repository of deep sequencing assemblies obtained with purified m-RNA from skate electroreceptors. This data could also be used to calculate expression levels, which cannot be appreciated using PCR alone. Since the calcium activated potassium channel (*Kcnma1*) and the voltage dependent calcium channel (Cav1.3) have already been characterized in this way, we turned our attention to the beta subunits of the BK channel, which affect gating properties, and to the fast voltage dependent potassium current (presumably an A current), which mediates the oscillatory responses of the ampulla that are thought to occur in free living skates. The deep sequencing data also tended to confirm our previous impression [21] that *Kcnmb1* is not expressed in the electroreceptor, and may not be present in the skate at all, given the absence of genomic contigs.

A complete assembly for skate *Kcnmb2* was found, and the sequence was confirmed by PCR and cloning. It was somewhat surprising to find that the expression levels for *Kcnmb2* were 1000 fold lower than for *Kcnma1*, which tends to suggest that this subunit plays no physiological role. Skate BK channels are thought to mediate accommodation of the electrorceptor to sustained potentials in the environment and to keep the receptor cells poised at threshold [9]. Rapid ball and chain inactivation of the skate BK channels by *Kcnmb2* would not be appropriate for this purpose, and there would be no such inactivation if the α subunits are indeed expressed at a 1000-fold higher level. In addition, skate *Kcnmb2* was found to have 51 additional amino acids upstream of the usual n-terminus, where the 45 amino acids that comprise the ball and chain are located [32]. It is postulated that these amino acids could prevent the occurrence of inactivation. Since the skate is capable of expressing *Kcnmb2* in the electroreceptor, it seems likely that other tissues, especially certain brain regions [19] would express β2 at levels that could have a physiological effect. We also note that the ratio of *Kcnma1* to β2 at the protein level, and in the tetrameric channels that reach the cell membrane could be much less than 1000 to one. BK channels are known to be present in mitochondria, and BK channels in mitochondria have a specific insert at alternative splice site 7 which controls targeting [33]. More recently, Li et al [34] have reported that BK channels are present in the nuclear envelope of hippocampal neurons, where they regulate membrane potential and modulate release of calcium into the nucleoplasm, which leads to expression of specific genes.

In the case of β4 the extremely low expression levels may explain why a complete sequence could not be obtained. Deep sequencing produced only one assembly containing β4 and that assembly contained only the first exon. We had anticipated that deep sequencing might succeed since multiple assemblies might be obtained, as seen with Kv1.1.

### Physiological Role of Kv1.1 in elasmobranch electroreceptors

Kv1.1 is a commonly expressed low threshold voltage dependent potassium channel of the Shaker family [35]. It is most often expressed in axons, although it is also found in the soma of neurons [36, 37]. Confocal microscopy of sensory neurons stained with a fluorescent antibody showed localization of Kv1.1 in the surface membrane with some diffuse staining of the cytoplasm, possibly the endoplasmic reticulum [36]. The biological samples described in this report included 2-5 millimeters of myelinated nerve attached to each ampulla. Actively translating mRNA has been described in embryonic peripheral axons in cell culture and embryonic peripheral axons in vivo [38]. However mRNA has not been found in myelinated peripheral nerves of mature vertebrates. Based on the physiological and pharmacological evidence for a voltage dependent K current in the basal membranes of the receptor cells [9] it is likely that the Kv1.1 and/or Kv1.5 transcripts found in this study give rise to K channels that carry this current. However, the exact subcellular location of the Kv1.1 and Kv1.5 transcripts, and the extent to which they colocalize with each other remain to be determined.

In this report, two assemblies were found in the deep sequencing data which represent the full length beta subunit of Kv1.1. These have been designated as skate *Kcnab2* based on their alignment with the human sequence (Fig. 6) and with many other species as can be seen using Blastp. The properties of *Kcnab2* have been described by McCormack et al [41]. *Kcnab2* (also called Kvβ2) is by far the most abundantly expressed Kv1 beta subunit in the brain. KV1.1 channels containing only the *Kcnab2* subunit do not exhibit rapid “ball and chain” inactivetion because the requisite amino acids at the N-terminus that mediate this inactivation are missing [41]. Kvβ1 and Kvβ3 have additional amino acids and do produce rapid inactivation when they are present. This can be shown by aligning the sequence of skate Kvβ2 in Fig. 6 with a human long form of human Kvβ1, NP_751892.1, which contains 419 amino acids. As noted above, the region of alignment begins at amino acid 77 of human Kvβ1 which means that the first 76 n-terminal AA’s of human Kvβ1 are missing in the skate. This substantiates the conclusion that a skate Kv1.1 channel that contained only Kvβ2 would not inactivate.

### Alignment of Elasmobranch *Kcnmb2* and *Kcnmb4* to the Lamprey, *P. marinus*

Lampreys are among the few species as ancient or more ancient than cartilaginous fish. Lampreys are of particular interest because they have electroreceptors. Recordings from lamprey brain areas known to mediate electroreception show responses to very weak electrical stimuli, comparable to those detected by elasmobranchs [42]. Lampreys have discrete receptors called “end buds.” End buds are similar to skate electroreceptors insofar as they have goblet shaped receptor cells which synapse onto an afferent fiber and are separated by supporting cells. Short lines or clusters of 2-8 end buds are distributed over the trunk or head of the animal [44]. Although the lampreys do not have ampullae or high resistance jelly filled canals, the evolutionary chart in Fig. 13 suggests that the common “ancestral vertebrate” (550 million years ago) may have had electroreceptors. The amino acid sequence of the beta two subunit of *Petromyzon marinus* is shown in the top line of Fig. 9 (Uniprot S4RHH3), and shows significant homology with the other species, with the exception that the additional amino acids found in the two elasmobranch species do not seem to be present in the lamprey. There is no data on expression levels of lamprey BK channels nor is it proven that BK is specifically present in end buds. However, it is entirely possible β2 and β4 both have a physiological role of the BK channel in the lamprey.

**Fig 13.**
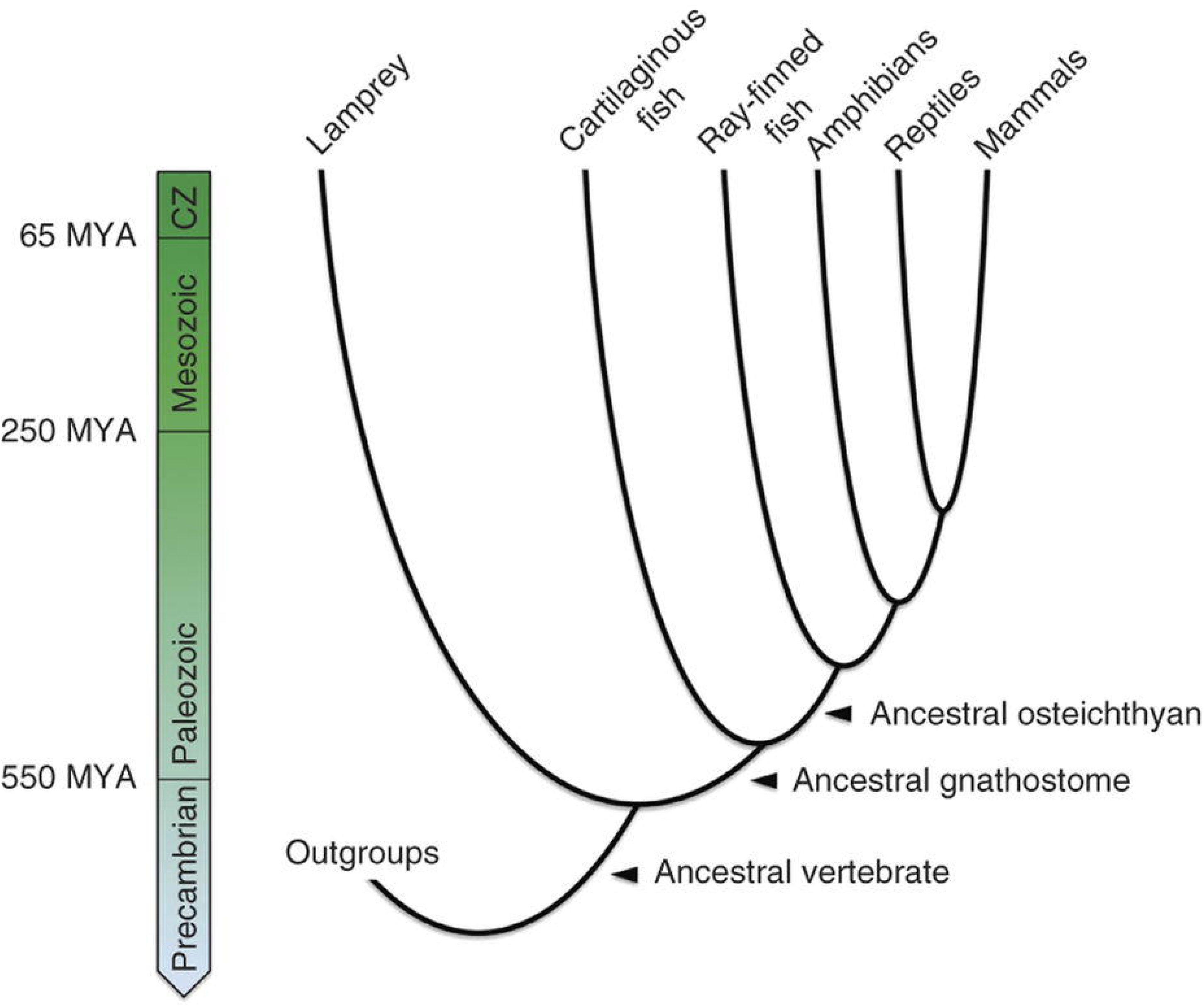
Abridged Phylogeny of the Vertebrates. Colecanths, lungfish, hagfish and certain extinct lineages have been excluded for simplicity. Cartilaginous fish include elasmobranchs (*R. typus* and *L. erinacea*) and *C.* milii a chimera. From ref. 44 with permission.

*P. marinus* also has a beta 4 subunit (Uniprot S4R636) which is strongly aligned with human beta 4 (NP_852006.1; E = 6e-43). *P marinus* beta 4 shows a similar degree of alignment with the assembled *L. erinacea* beta 4 [21] with E = 8e-42. This observation is not surprising because the beta 4 subunit shows much greater conservation across species than beta 2 (see Fig. 9 of ref 21). Based on the diagram in Fig. 13, it can be concluded that the “common ancestral vertebrate” is likely to have had an orotholog of the genes for both beta 2 and beta 4, and that these genes functioned in the nervous system, which probably included electroreceptors.

### Probable evolutionary origin of the Beta Subunits of *Kcnma1*

Higher vertebrates including humans are known to have four beta subunits, *Kcnmb1-4*, which seem to have arisen through a series of gene duplications. This can be inferred from the degree to which the human sequences for *Kcnmb1, Kcnmb3* and *Kcnmb4* can be aligned with *Kcnmb2*. *Kcnmb2* has 235 amino acids in humans. The number of amino acids in the other subunits, and the E-value for alignment against human *Kcnmb2* are as follow: *Kcnmb1*: 191 aa, 7e-56; *Kcnmb3*: 277 aa, 7e-65; *Kcnmb4*: 210 aa, 3e-39. Several gene duplications would have been necessary to explain the four orthologs, and cross species comparisons shed insight on when these duplications occurred. There are two important clues. First, as noted above, *Kcnmb2* and *Kcnmb4* seem to be ubiquitous, and are likely to have been present in the ancestral vertebrate (Fig. 13). Of these, *Kcnmb4* shows the least variation among species. The second clue is that *Kcnmb1* is not present in *P. marinus* or in ray finned fish (Actinopterygii) or in any cartilaginous fish. As previously reported, we have searched for *Kcnmb1* in Skatebase by entering the human nucleotide sequence and no significant alignment with genomic or transcriptome contigs was found (21). Attempts to locate *Kcnmb1* in the transcriptome of *L. erinacea* ampulla of Lorenzini (deep sequencing data [29] and genomic contigs) yielded only *Kcnmb2*. Searches using BlastP have also failed to find an ortholog of *Kcnmb1* in either *c. milii or r. typus.* This suggests that *Kcnmb1* first appeared in lobe finned fish. The most basal lobe finned fish with a genome sequence and electroreceptors is the coelacanth, *Latimeria chalumunea* [45]; and it has a *Kcnmb1* gene, as well as genes *for Kcnmb2-4*. However, entry of the human nuceleotides for *Kcnmb3* into Skatebase gives no significant alignments to any genomic or transcriptome contig. Moreover a search of annotated databases for possible orthologs of *Kcnmb3* in *L. erinacea, c. milii, r. typus and P marinus* also does not reveal any. Based on Fig. 13 and the similarity of the coelacath’s fleshy fins to legs, the coelacanth, is thought to have existed unchanged for about two hundred million years [38], and to closely resemble ancestors of earliest amphibians. It is concluded that all of the gene duplication events that lead to the existence four beta subunits probably occurred before there were land dwelling vertebrates, but that catilagenous fish may have had only *Kcnmb2* and *Kcnmb4*.

### Future Directions

Additional studies should include expression of the BK channel, and the newly discovered Shaker channels in oocytes followed by single channel recordings. Physiological studies of the short Kv1.5 are particularly important because this is a newly discovered and entirely novel channel based on the length of the alpha subunit. Studies should include both Kv1.1 and Kv1.5 in the presence and absence of the beta subunit, *Kcnab2*. Quantitative PCR studies should be performed to determine the relative expression levels of Kv1.1 and Kv1.5. Immunofluorescence studies should also be performed since it is not known whether Kv1.1 and Kv1.5 co-localize with one another at the cellular level. Commercially available antibodies are available whose immunogens show homology with the sequences of skate Kv1.1 and Kv1.5. Studies involving these antibodies are expected to confirm protein expression at the basal faces of the receptor cells. Studies of the *Kcnmb2* subunit of BK should include Western blots, which might be used to confirm the presence of additional amino acids at the N-terminus. Experiments should be done in skate brain, parts of which may have higher expression of *Kcnmb2* than the electroreceptor. It would also be useful to determine whether the newly discovered short Kv1.5 sequence is expressed in the skate brain, and which regions.

## Acknowledgement

We are very grateful to Dr. Benjamin King for important assistance with informatics. We also thank Clare Baker of Cambridge University for helpful comments and suggestions.

